# Insular input to the prelimbic cortex underlies social affective behavior in rats

**DOI:** 10.1101/2025.10.02.680023

**Authors:** E.K. Chun, A. Djerdjaj, T. Matulis, A.J. Ng, M.M. Rogers-Carter, J.P. Christianson

**Author notes:** corresponding Author: John Christianson, 1-617-552-3979. denotes equal contributions (E.K. Chun, A. Djerdjaj).

## Abstract

To navigate social interactions, animals must adjust their behavior in response to information derived from conspecifics. The integration of social information and coordination of behavior occurs within a distributed social decision making network. The prelimbic (PL) prefrontal cortex and the insula (IC) are key nodes in the salience network which is anatomically situated to interact with the social brain. We investigated the IC-PL circuit in a social affective preference (SAP) test in which subject rats are exposed to 2 age-matched conspecifics where one is stressed via footshock and the other is naive to stress. Typically, rats approach stressed juvenile conspecifics but avoid stressed adults. Using a combination of local and tract specific loss of function methods, we demonstrate that the PL, anterior IC, and the tracts between the posterior or anterior IC and the PL are necessary when rats face the choice to approach or avoid stressed conspecifics. Going further, chemogenetic inhibition of PL neurons innervated by the IC also interfered with social affective behaviors. These studies enrich our understanding of the neurobiology of social decision making by establishing a mechanistic link between insular and prefrontal circuits.

**Significance Statement:** The successful navigation of social interactions requires detection of the emotional states of others and the appropriate behavioral output. While the insula and prefrontal cortex have both been implicated as key brain regions in social decision-making and the salience network, their functional role in guiding social behaviors are largely unknown. Using a combination of region-specific and circuit manipulations in rats, we present evidence of the necessity of the insula-prefrontal pathway in the approach and avoidance of stressed others, reinforcing the importance of this cortical system in social affective processing. This work provides new insight into the network-level mechanisms underlying social behavior and highlights an important circuit that may be relevant to understanding neuropsychiatric disorders with social impairments.

## Introduction

Social interactions are the product of complex, multilevel information processing and decision making. As such, social cognition is thought to emerge from a distributed set of brain networks that provide for the detection and appraisal of social and contextual information leading to the organization of situationally appropriate behaviors (Newman, 1999; Goodson, 2005; O’Connell and Hofmann, 2011; Prounis and Ophir, 2020). Cortical circuits are implicated in a wide range of social and affective processes and there is increasing emphasis on understanding how the intrinsic functional networks of the brain interact to guide behavior (Uddin, 2015). The salience network consisting of the insula and frontal cortex is engaged by a variety of stimuli related to interoceptive and homeostatic needs and, naturally, social functions are correlated with activity in the salience network (Seeley, 2019). Thus, dysfunction in the salience network may underlie a wide range of psychopathologies (Menon, 2011; Goodkind et al., 2015), including those with severe social abnormalities (Uddin et al., 2013; Guo et al., 2019; Tavares et al., 2022).

Rodents appear to have an homologous salience network that includes insular cortex and regions of medial prefrontal cortex (Tsai et al., 2020). The prefrontal cortex is involved in social learning, social recognition and other aspects of social behavior (Liang et al., 2018; Levy et al., 2019; Xu et al., 2019; Frost et al., 2021) and the insular cortex is involved in approach/avoidance behaviors to novel, sick, and distressed conspecifics (Rogers-Carter et al., 2018b; Miura et al., 2020; Kim et al., 2022; Rieger et al., 2022b). Because the insula is involved in social affective and sensory processes, and the prefrontal cortex is involved in organizing goal-directed behaviors, we hypothesized that rodent social affective behaviors would require signalling in the insula-prefrontal circuit. We focused on the prelimbic (PL) region because it receives input from the extent of the insula (Hoover and Vertes, 2007), encodes social novelty (Zhao et al., 2022) among other functions in the social realm, and has projections to downstream regions that, when altered, contribute to anxiety- and autism-related behaviors (Luo et al., 2023). Additionally, a network analysis revealed the PL to be the strongest participant in the brain network response to stressed conspecifics (Rogers-Carter et al., 2018b), potentially positioning it as a crucial node necessary for social decision-making.

In our prior work, we established that the posterior insular cortex (pIC) is necessary and sufficient for social approach to stressed juvenile conspecifics and social avoidance of stressed adults (Rogers-Carter et al., 2018b, 2019; Rieger et al., 2022a). The pIC innervates both the anterior insula (aIC) and PL (Shi and Cassell, 1998; Hoover and Vertes, 2007; Gehrlach et al., 2020) so here we used a combination of pharmacology, tract tracing, and circuit chemogenetics to establish what the aIC, PL, and the inputs from the aIC and pIC to the PL contribute to rat social affective behaviors.

We first tested the PL’s role in SAP testing by infusing muscimol to the region. We then used retrograde tracing to map insular inputs to the PL and found that both the aIC and pIC provide strong input into the PL. Because we had not previously tested the role of the aIC itself in the SAP paradigm, we next used chemogenetic inhibition of the aIC in SAP tests finding that it, like pIC, is also involved in social approach decisions. To determine the role of the inputs of the insula to the PL, we took two approaches. We used chemogenetics to inhibit insula axon terminals within the PL and we used a transsynaptic approach to inhibit PL neurons that receive insula input.

Finally, we stained the insula-innervated PL neurons to describe the distribution of glutamatergic or GABAergic neurons in this circuit. In sum, we report that this insula-PL pathway is necessary for social affective behaviors towards stressed others positioning this circuit as critical to social contexts where affective discrimination is required.

## Materials and Methods

### Animals

Male Sprague-Dawley rats were purchased from Charles River Laboratories (Wilmington, MA) and allowed to acclimate to the vivarium in the Boston College Animal Care Facility for at least 7 days before any treatment. Adult experimental rats arrived weighing 225-250g, juvenile conspecifics arrived at PN21, and adult conspecifics arrived at PN55. Experimental rats were housed in pairs while juvenile and adult conspecifics were housed in triads. The vivarium maintained a 12h light/dark cycle and food and water were available *ad libitum*. Behavioral testing was conducted within the first 4 hours of the light cycle. All procedures were approved by the Boston College Institution Animal Care and Use Committee and adhered to the Public Health Service *Guide for the Care and Use of Laboratory Animals*.

### Surgical procedures - Retrograde tracers

To quantify insula projections to the PL, Fluorogold (4%, Fluorochrome), a retrograde tracer, was deposited unilaterally into the PL. Adult male rats underwent surgery under inhaled anesthesia (2-5% v/v isoflurane in O_2_). Fluorogold was unilaterally microinjected into the PL (from bregma: A/P +3.2, M/L +/-0.6 D/V -3.5) at a rate of 100nL/min to a total volume of 100nL with 5 min allowed for diffusion. After surgery, rats were given prophylactic meloxicam (1 mg/kg, s.c., Eloxiject; Henry Schein) and penicillin (1 mg/kg, s.c., Combi-Pen; Henry Schein). 10 mL lactated Ringer’s solution was delivered in two doses of 5 mL (s.c.).

### Surgical Procedures - Cannula Implantation

To inhibit PL activity, indwelling cannula were implanted bilaterally into the PL to allow for direct infusion of muscimol or vehicle. Experimental adult males underwent surgery under anesthesia as above. Bilateral cannula (26g, Plastics One) were implanted into the PL (from bregma: A/P +3.0, M/L +/-0.5 D/V -3.2). Cannula were fixed in place with stainless steel screws and acrylic cement and were fitted with stylets to maintain patency. Postoperative care was as described for fluorogold. Rats were allowed 3 weeks of recovery before undergoing the behavioral procedures described below. 1 week prior to behavioral testing, test rats were wrapped in a cotton towel and handled by an experimenter for 1 minute each weekday for a total of 5 days to habituate rats to handling prior to drug infusion. To inhibit the PL, 1 h prior to SAP testing, muscimol (100 ng/side in 0.5mL of saline, Tocris) was microinjected at a rate of 1μL/min with an additional 1 minute diffusion time as previously (Chen et al., 2016; Rogers-Carter et al., 2018b; Sarlitto et al., 2018). After behavioral testing was finished, experimental rats were overdosed with tribromoethanol and brains were removed, flash frozen, and sectioned at 40 μm using a freezing cryostat (Leica CM1860 UV), mounted on gelatin-coated slides, and stained with cresyl violet to locate cannula placements.

### Surgical procedures - Chemogenetic manipulations

We utilized viral transfer of the chemogenetic Gi coupled receptor hM4Di (Rogan and Roth, 2011) to manipulate the activity of the aIC, the inputs of the aIC and pIC to the PL, and the neurons of the PL that are postsynaptic to aIC or pIC afferents. We used a viral vector that targets neurons (AAV5-hSyn-hM4Di-mCherry; Addgene Cat.

No. 50475-AAV5; titer = 8.6x10^12^ GC/mL or Cre dependent AAV5-hSyn-DIO-hM4Di; Addgene Cat. No. 44362-AAV5; titer=2.4x10^13^ GC/mL). To inhibit the aIC, 500nL of hM4Di was deposited bilaterally. To inhibit the aIC or pIC axon terminals in the PL, hM4Di was delivered bilaterally to either the aIC or pIC and a bilateral cannula was placed in the PL (exactly as described for muscimol injections) for later microinfusion of the hM4Di agonist deschloroclozapine (DCZ, Tocris). The coordinates for aIC deposits were from bregma: A/P +2.7, M/L +/-4.3 D/V -5.6 and for pIC from bregma: A/P -1.8, M/L +/6.5 D/V -7.

We previously validated the specificity and efficacy of these viruses in the insula-nucleus accumbens tract (Rogers-Carter et al., 2019) and basolateral amygdala-insula tract (Djerdjaj et al., 2022) using the chemogenetic ligand, clozapine-n-oxide (CNO). In the present set of experiments, we used CNO when targeting prelimbic neurons and DCZ in the other applications. Although presented last in the report, data collection for the PL_IC_ experiments began before the publication of work describing greater efficacy of DCZ (Nagai et al., 2020; Nentwig et al., 2022) which prompted us to switch to DCZ for the ensuing experiments. We verified the functional utility of DCZ in all of these preparations (Supplementary Figure S1).

To target the PL neurons postsynaptic to insula input (PL_IC_ neurons), an anterograde, transsynaptic AAV encoding Cre-recombinase (AAV1-hSyn-Cre-WPRE-hGH; Addgene Cat. No. 105553-AAV1; titer=1.9x10^13^ GC/mL; hereafter: “AAV1-Cre”) was bilaterally injected into either the aIC or pIC and the Cre-dependent hM4Di vector was bilaterally injected into the PL (from bregma: A/P +3.2, M/L +/-0.6 D/V -3.5). All virus infusion surgeries were carried out under inhaled anesthesia with postoperative care as described above. The viral approach taken here transfers Cre-recombinase to all monosynaptic target cells of the insula and, in this case, induces expression of hM4Di receptors in these cells in the PL (Zingg et al., 2017, 2020), allowing for reversible inhibition of putative PL_aIC_ and PL_pIC_ neurons via systemic administration of the hM4Di actuator, CNO (Rogan and Roth, 2011). Rats were allowed 3 weeks of recovery before undergoing behavioral procedures as described below. To inhibit these specific cell populations, rats received either an intraperitoneal (I.P.) vehicle (DMSO in Saline) or CNO (3 mg/kg; Tocris) injection 45 minutes prior to SAP testing.

### Social Affective Preference (SAP) Tests

This procedure allows for the quantification of a rodent’s behavior when interacting with conspecifics of varying social affect. The SAP tests were conducted in plastic arenas (76.2cm × 20.3cm × 17.8 cm, L × W × H) with beta chip bedding and a transparent plastic lid. Conspecifics were placed into individual clear acrylic chambers (18 x 21 x 10cm; L x W x H) with walls made of acrylic rods spaced 1 cm apart that allowed for social interaction. During testing, the chambers containing conspecifics were placed at opposite ends of the arena. Testing typically consisted of 2 habituation days followed by up to 6 test days (*note that some experiments used modifications of this approach as noted in the results section). On day 1 (arena habituation), experimental rats were placed in the testing room for 1h and exposed to the testing arena for 15 minutes before being returned to their home cage. On day 2 (social habituation), experimental rats were placed in the plastic testing arena for 5 min of behavior testing where they were presented with 2 naïve juvenile conspecifics. On days 3 and 4, we tested the experimental rats’ social affective preference with juvenile conspecifics, on days 5 and 6 with adult conspecifics. Some experiments included 2 additional days on which we tested opposite sex preference (see Supplemental Figure S2). Tests occurred 45 minutes after administration of drugs (muscimol, CNO, or DCZ). For each pair of test days, the order of vehicle or drug injection was counterbalanced.

On each SAP test day, rats were transferred to the test room 1 h before testing. Tests began by transferring a rat to the test arena which contained a pair of unfamiliar conspecifics. To manipulate conspecific affect, one of the conspecifics received 2 footshocks immediately preceding placement in its chamber of the testing arena (5s, 1 mA, inter-shock interval of 50s); the other conspecific was naïve to any treatment. A trained observer quantified the amount of time the experimental rat spent investigating each conspecific. Social investigation was defined as time spent sniffing or touching the conspecific through the acrylic bars. A second trained observer, blind to experimental conditions, scored digital videos to establish inter-rater reliability. With the exception of the muscimol experiment, all testing followed a counterbalanced, within-subjects design.

### Tissue collection

Rats from the chemogenetic studies were overdosed with tribromoethanol and perfused with cold 0.01 M heparinized PBS followed by 4% paraformaldehyde.

Dissected brains were stored in 4% paraformaldehyde for 24 h and then transferred to 30% sucrose for at least 2 days. 40 µm coronal sections were obtained in series on a freezing cryostat. For the tracer study, PL and insula sections from along the rostral-caudal axis were directly mounted onto slides and coverslipped with Vectashield containing DAPI (Vector Laboratories) to visualize Fluorogold in the PL for verification, and the anterior, medial, and posterior insula for quantification of projection patterns. For the chemogenetic studies, aIC, pIC, and PL sections were directly mounted onto slides and coverslipped with Vectashield containing DAPI to visualize mCherry, a genetically encoded fluorescent protein fused to hM4Di, under a fluorescent microscope to determine the spread of hM4Di. Additional sections containing either anterior or posterior insula were collected and stored in cryoprotectant for later verification of cre-recombinase with immunohistochemistry.

### Immunohistochemistry

To verify expression of cre recombinase in the anterior or posterior insula, tissue sections were washed in PBS-T (0.01% Triton-X 100), blocked in 5% normal donkey serum in PBS-T, and then incubated overnight in mouse anti-cre recombinase primary antibody (1:5,000; EMD Millipore, Cat. No. MAB3120). Sections were then washed in PBS-T and incubated in AlexaFluor 488 AffiniPure donkey anti-mouse fluorescent secondary antibody (1:500; Jackson Immunoresearch, Cat. No. 715-545-150). Sections were then floated onto glass slides and coverslipped with Vectashield containing DAPI. To identify and quantify PL_IC_ cell-type as glutamatergic or GABAergic, sections from rats with AAV1-Cre and mCherry in the PL were stained for either CaMKii or GAD67.

Sections were washed in PBS-T and blocked in 5% normal donkey serum. One set of tissue (n=30) was incubated overnight in mouse anti-CaMKII alpha primary antibody (1:5,000; Invitrogen, Cat. No. MA1-048) and a second set of tissue (n=30) in mouse anti-GAD67 primary antibody (1:5,000; Sigma, Cat. No. MAD5406). Sections were washed in PBS-T and incubated in AlexFluor 647 AffiniPure donkey anti-mouse fluorescent secondary antibody (1:500; Jackson Immunoresearch, Cat. No. 15-605-150). Sections were then floated onto glass slides and coverslipped with Vectashield containing DAPI.

### Imaging and cell-type quantification

All imaging was conducted with a Zeiss Axioimager Z.2 fluorescent microscope in the Boston College Imaging Core. Z-stacked, tiled images containing the ROIs (anterior, medial, or posterior insula for retrograde tracing study; PL for neuroanatomy study) were taken at 20x using a Hammamatsu Orca Flash 4.0 digital camera. Using ImageJ software, individual fluorescent channel images were stacked, ROIs were traced with reference to the rat brain atlas, and the cell counter plug-in was used to count labeled cells. For the retrograde tracing study, cells that expressed Fluorogold were quantified and expressed as a percentage of DAPI cells. For the neuroanatomy study, superficial (layers II/III) and deep (layers V/VI) ROIs of the PL were counted within each image. For each ROI, the total number of DAPI, mCherry-expressing PL_IC_ cells, CaMKII or GAD67 cells, and PL_IC_ cells coexpressing either CaMKII or GAD67 was quantified within this population to determine whether PL_IC_ cells were primarily glutamatergic or GABAergic.

### Experimental Design and Statistical Analysis

Data from rats receiving stereotaxic injections were only included when site-specificity criteria were met. For muscimol studies, rats were included if cannula terminated within the PL. For chemogenetic studies, inclusion required both mCherry expression in the aIC or PL and Cre recombinase expression in the anterior or posterior insula. These strict parameters necessitated the exclusion of a number of rats within each cohort. Due to this, and to minimize the number of animals used, some rats with unilateral viral expression in the PL and insula were included. Sample sizes were determined by conducting a priori power analyses in G*Power using effect sizes observed previously (Rogers-Carter, 2018) and N = 12 was determined to be appropriate to achieve power ≥ 0.70. Datasets were tested for normality and sphericity prior to analysis and found to be suitable for t-test and analysis of variance (ANOVA).

For the retrograde tracing, fluorogold expression was variable across subjects despite consistent targeting of the PL. Due to this study being confirmatory rather than descriptive in nature, data were analyzed from one representative rat that displayed Fluorogold expression in the PL and robust retrograde expression along the insula. An one-way ANOVA was used to compare fluorogold and DAPI co-labeled cells across insular subregions (anterior vs. medial vs. posterior). This method was also used to compare co-labeled cells as a percentage of DAPI cells. Post-hoc analysis consisted of Tukey’s multiple comparisons test. For muscimol experiments, data were analyzed using a between-subjects two-way ANOVA. For chemogenetic experiments, social exploration times were compared using repeated measures two-way ANOVAs with conspecific stress and drug treatment (vehicle or CNO/DCZ) as within-subjects variables. Main effects and interaction effects were deemed significant at p < 0.05 and followed by Sidak post-hoc tests to maintain experiment-wise type 1 error rate to ɑ < 0.05. To control for individual variability in social interaction time, preference for the stressed conspecific in each condition was calculated as a percentage of the total time spent investigating both conspecifics (time interacting with stressed / (time interacting with naïve + time interacting with stressed) x 100). In muscimol experiments, preference scores were compared using an unpaired t-test. In chemogenetic experiments, preference scores were compared using repeated-measures two-way ANOVA with age (juvenile or adult) and drug (vehicle or CNO) as within-subjects factors. Two-way ANOVA was used to analyze differences in PL_IC_ population size and cell-type distribution across cortical layers (layer II/III vs. layers V/VI) and between insula projection origin (anterior vs. posterior). Three-way ANOVA was used to determine the primary cell-type of PL_IC_ neurons (GAD67 vs. CaMKII) and whether this varied based on cortical layer (layer II/III vs. layers V/VI) or insula projection origin (anterior vs. posterior). P values and the effect size (𝜂^2^ or partial 𝜂^2^) are provided for significant main effects and interactions. Statistical analyses were conducted in Prism 10.6 (Graphpad Software).

## Results

### The PL is necessary for social affective preference behavior toward stressed juveniles

To determine the effect of PL inactivation on social affective behavior, adult male rats received bilateral cannula implants in the PL and later underwent SAP testing with juvenile conspecifics after vehicle or muscimol injections (Figure 1A-D). After cannula verifications, 22 rats met inclusion criteria. Experimental rats in the vehicle condition spent more time investigating stressed juveniles while rats that received muscimol injections had no preference (Figure 1B). There was a drug by affect interaction (*F*(1,20) = 12.0, p = 0.0025, 𝜂^2^ = 0.17) and a significant difference between social investigation of naïve and stressed juveniles in the vehicle condition (p = 0.004) that was not present in the muscimol condition (p = 0.376). The preference for the stressed conspecific was calculated as a percentage of the total time spent investigating both conspecifics (Figure 1C), which revealed a significant preference for the stressed juvenile in the vehicle rats compared to the rats that received muscimol (t(20)= 3.52, p = 0.002). In sum, inactivation of the PL via muscimol administration interfered with a rat’s preference for stressed juveniles in the SAP test.

**Figure 1.**
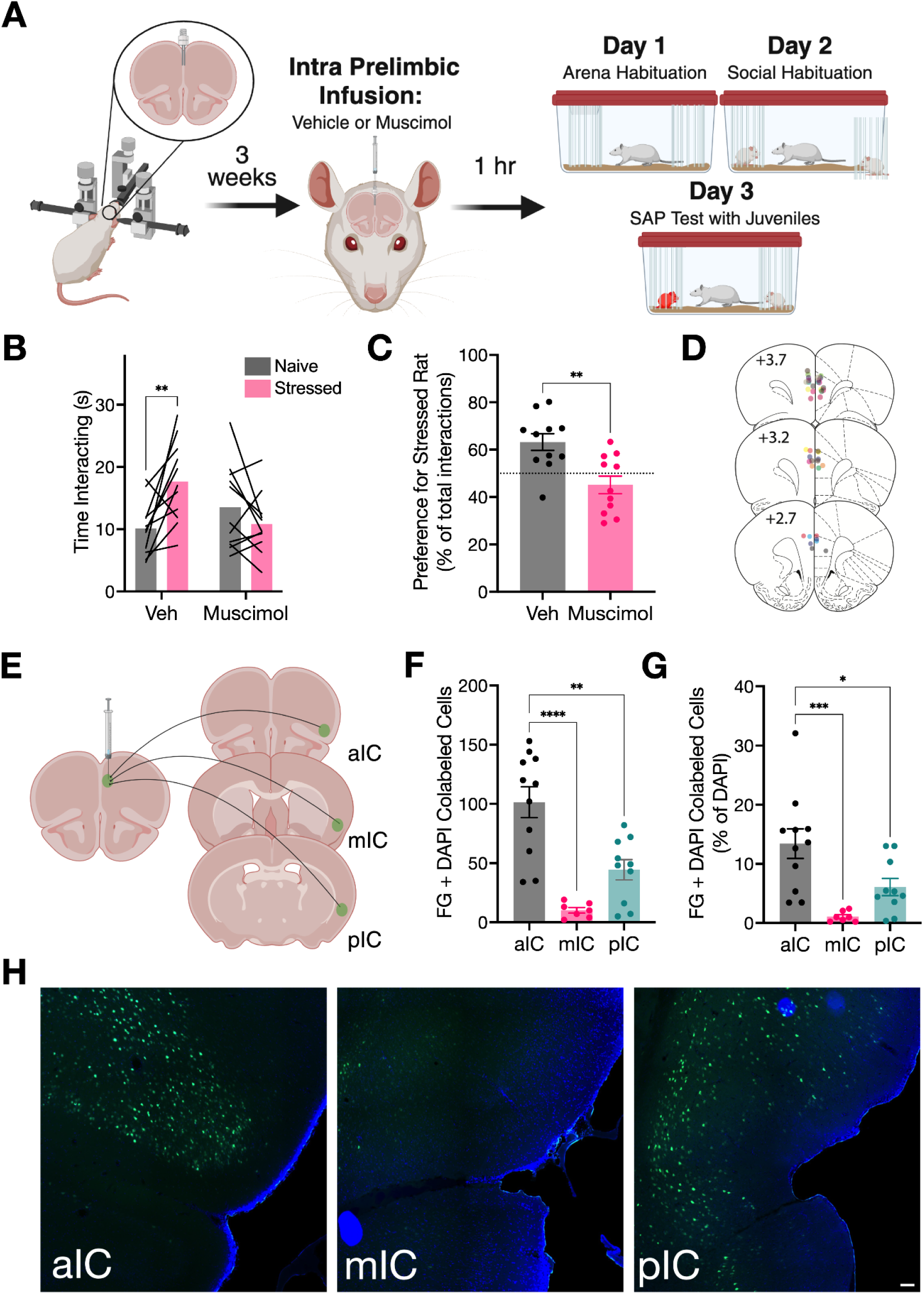
PL inactivation abolishes preference for stressed juveniles. **A.** Schematic diagram of the experimental design. Bilateral cannula were implanted into the PL of male test rats and, 3 weeks later, rats received vehicle (Veh) or muscimol injections one hour prior to SAP tests with juvenile conspecifics. **B.** Mean (with individual replicates) time spent exploring the naïve and stressed conspecifics during the 5 min SAP test. Rats in the vehicle condition preferred social investigation of stressed juveniles compared to naïve juveniles (p = 0.004), while rats that received muscimol injections showed no difference in social investigation. **C.** Data from **B** presented as a preference for the stressed conspecific as a percentage of total social interaction. Experimental rats in the vehicle condition preferred interaction with stressed juveniles, while rats in the muscimol condition displayed no preference (p = 0.002). **D.** Location of cannula implants in the PL (individual rats are represented by different colors). Atlas images recreated from Paxinos & Watson (1998), use pending permission. **E.** Schematic of experimental design. Fluorogold was unilaterally deposited into the PL of male test rats. Retrograde expression of fluorogold was quantified along the rostral-caudal gradient of the insula. **F.** Mean (+/- SEM) number of colabeled cells (fluorogold + DAPI). Each dot represents data from one fluorescent image. The anterior portion of the insula had significantly more colabeled cells than both the medial (p < 0.0001) and posterior (p = 0.001) insula. **G.** Data from **F** expressed as a percentage of DAPI cells. Anterior insula had a higher percentage of PL-projecting cells than both the medial (p = 0.0006) and posterior (p = 0.024). **H.** Representative fluorescent images of anterior (aIC), medial (mIC), and posterior (pIC) insula (blue = DAPI, green = fluorogold). Scale bar = 200 μM. Diagrams in A & E created with BioRender.com. *p < 0.05, **p < 0.01, ***p < 0.001, ****p < 0.0001.

### Anterior and posterior subregions of the insula project to the PL

Insula projections to the medial prefrontal cortex, and to the PL specifically, are well-documented in rodent models (Vertes, 2004; Hoover and Vertes, 2007; Mathiasen et al., 2023). To confirm the extent of these projections in Sprague-Dawley rats, fluorogold, a retrograde tracer, was deposited into the PL of adult male rats (Figure 1 E-H). Ten days later, rats were perfused, brains were extracted, and tissue was collected for cell quantification. Cells expressing DAPI, a nuclear stain, and green fluorescence, which marked the location of flurogold in insula afferents to the PL, were counted in the anterior, medial, and posterior portions of the insula (Figure 1H). We observed the greatest number of fluorogold positive cells in the anterior and posterior insula, which resulted in a main effect of ROI in both the number of colabeled cells (*F*(2,25) = 18.9, p < 0.0001) and the percentage of colabeled cells (*F*(2,25) = 9.85, p = 0.0007), with anterior insula having significantly more projections to the PL than both the medial (p < 0.0001) and posterior (p = 0.0003) insula. In sum, both aIC and pIC project to the PL with a denser projection originating from the anterior insula.

### Anterior insula inactivation interferes with social affective preference

Several recent reports provide evidence that the aIC contributes to a range of social behaviors (Cox et al., 2022; Kim et al., 2022), however we have not previously investigated its role in social affective behavior. hM4Di was introduced to the aIC and we conducted juvenile and adult SAP tests after vehicle or DCZ injections (0.1 mg/kg i.p., Figure 2A). After vehicle injections, test rats spent more time investigating stressed juveniles but displayed no preference after DCZ (Figure 2C). A significant stress by drug interaction (*F*(1, 12)= 12.43, p = 0.004, 𝜂^2^ = 0.16) and significant posthoc comparisons between the stress and naive exploration times (p = 0.005). After vehicle injections, test rats spent less time interacting with stressed adults without a clear trend in rats treated with DCZ (Figure 2D). This pattern is supported by a significant main effect of drug, (*F*(1, 12) = 11.44, p = 0.005, 𝜂^2^ = 0.11), drug by stress interaction (*F*(1,12) = 4.764, p = 0.050, 𝜂^2^ = 0.06) and significant post hoc comparison between stress and naive in the vehicle condition (p = 0.006). The percent time spent interacting with stressed conspecifics (Figure 2E) shows a strong effect of age (*F*(1,12) = 13.44, p = 0.003, 𝜂^2^ = 0.19) and an age by drug interaction (*F*(1, 12) = 18.03, p = 0.001, 𝜂^2^ = 0.26). The strongest effect of aIC inhibition was present in juvenile preferences (p = 0.004) while the comparison between stress and drug in adults did not reach significance (p = 0.128). Together these results suggest that aIC inhibition disrupts social approach to stressed juveniles and to a lesser extent, avoidance of stressed adults.

**Figure 2.**
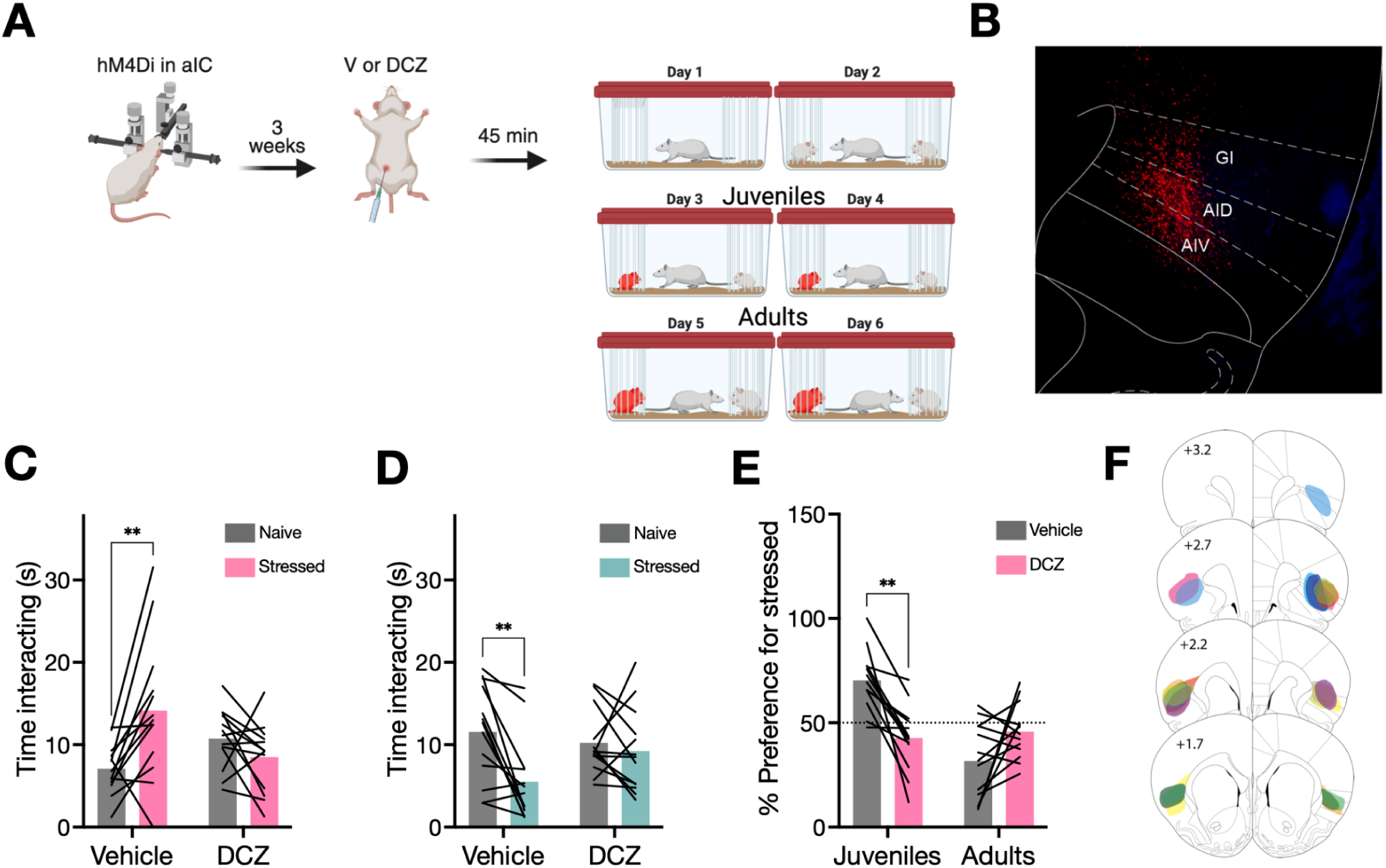
Anterior Insula inactivation interferes with social affective preference. **A.** Schematic diagram of the experimental design. Bilateral deposits of AAV5-hSyn-hM4Di-mCherry (hM4Di) were made in the aIC of male test rats and, 3 weeks later, rats received vehicle (V) or deschloroclozapine (DCZ) injections 45m prior to SAP tests with juvenile conspecifics and then adult conspecifics. **B.** Representative location of hM4Di deposit in aIC. **C.** Mean (with individual replicates) time spent exploring the naïve and stressed conspecifics during the 5 min SAP test. Rats in the vehicle condition preferred social investigation of stressed juveniles compared to naïve juveniles (p = 0.005), while rats that received DCZ injections showed no difference in social investigation. **D.** Mean (with individual replicates) time spent exploring the naïve and stressed conspecifics during the 5 min SAP test with adults. Rats in the vehicle condition avoided social investigation of stressed adults (p = 0.006), while rats that received DCZ injections showed no difference in social investigation. **E.** Data from **C** and **D** presented as a preference for the stressed conspecific as a percentage of total social interaction. Experimental rats in the vehicle condition preferred interaction with stressed juveniles and avoided stressed adults; DCZ eliminated the preference in juveniles (p = 0.004). **F.** Location of hM4Di expression in the aIC (individual rats are represented by different colors). Atlas images recreated from Paxinos & Watson (1998), use pending permission. Diagram in A created with BioRender.com.

### Terminal inhibition of insular afferents to the PL interferes with social affective preference

In our published and preceding work, we have established that the aIC and pIC innervate the PL and that isolated inhibition of the pIC, aIC or PL all interfere with social affective behaviors. To determine if input from the insula to the PL is necessary for social affective behaviors, we inhibited synaptic transmission at the insula-PL synapses during SAP tests. To this end, rats received hM4Di in either the aIC or pIC and cannula for direct infusion of DCZ (500nl of 30 nM DCZ) to the PL 45 min before SAP tests (Figure 3A-C). For rats with aIC-PL hM4Di, rats spent more time interacting with the stressed juvenile conspecific after vehicle injections, and less time with stressed rats after DCZ injections (Figure 3D). This was supported by a significant stress by drug interaction (*F*(1, 12) = 26.08, p = 0.0003, 𝜂^2^ = 0.16) and significant post hoc comparisons between stress and naive in vehicle (p = 0.0067) and DCZ conditions (p = 0.0076). In SAP tests with adults, vehicle-treated rats spent less time interacting with the stressed conspecific after vehicle injection and equal time between stressed and naive conspecifics after DCZ (Figure 3E). This was supported by a significant stress by drug interaction (*F*(1, 12) = 8.825, p = 0.0117, 𝜂^2^ = 0.06) and significant post hoc comparison between stress and naive in the vehicle condition (p = 0.010). When considering the percent of time spent with the stressed conspecific, it was clear that DCZ treatment strongly influenced preference for the stressed juvenile and eliminated the avoidance of the stressed adults (Figure 3F) which was supported by a significant age by drug interaction (*F*(1, 24) = 35.20, p < 0.001, 𝜂^2^ = 0.29) and significant post hoc comparisons between vehicle and drug in juvenile (p < 0.001) and adult (p = 0.009) tests.

**Figure 3.**
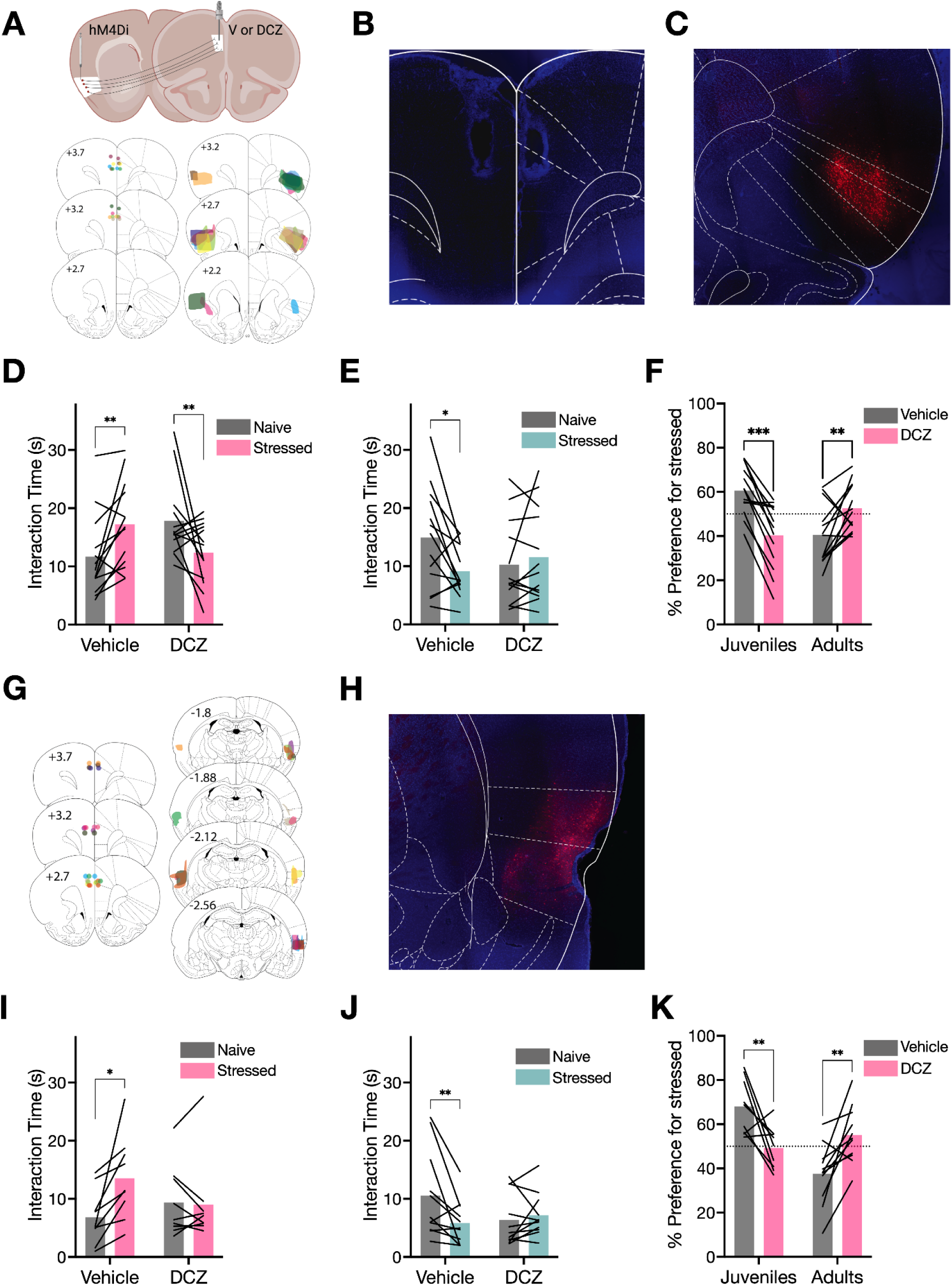
Terminal inhibition of insular afferents to the PL interferes with social affective preference. **A.** Schematic diagram of experimental design. AAV5-hSyn-hM4Di-mcherry (hM4Di) was deposited bilaterally in either the anterior or posterior insula (aIC or pIC) and bilateral cannula were implanted in the PL for delivery of vehicle or DCZ prior to SAP tests. Atlas images depict location of PL cannula and aIC virus placements corresponding to data in panels D-F. **B.** Representative image of PL cannula placement. **C.** Representative image of aIC hM4Di expression. **D.** Mean (with individual replicates) time spent exploring the naïve and stressed juvenile conspecifics during the 5 min SAP test. Rats in the vehicle condition preferred social investigation of stressed juveniles compared to naïve juveniles (p = 0.0067), while rats that received DCZ injections showed the opposite pattern (p = 0.0076). **E.** Mean (with individual replicates) time spent exploring the naïve and stressed conspecifics during the 5 min SAP test with adults. Rats in the vehicle condition avoided social investigation of stressed adults (p = 0.010), while rats that received DCZ injections showed no difference in social investigation. **F.** Data from **D** and **E** presented as a preference for the stressed conspecific as a percentage of total social interaction. Experimental rats in the vehicle condition preferred interaction with stressed juveniles and avoided stressed adults (p < 0.001); DCZ eliminated these preferences (p = 0.009). **G.** Location of PL cannula placements and pIC virus expression corresponding to data in panels I-K. **H.** Representative image of pIC hM4Di expression. **I.** Mean (with individual replicates) time spent exploring the naïve and stressed juvenile conspecifics during the 5 min SAP test. Rats in the vehicle condition preferred social investigation of stressed juveniles compared to naïve juveniles (p = 0.011), while rats that received DCZ injections showed no difference in social investigation. **J.** Mean (with individual replicates) time spent exploring the naïve and stressed conspecifics during the 5 min SAP test with adults. Rats in the vehicle condition avoided social investigation of stressed adults (p = 0.0038), while rats that received DCZ injections showed no difference in social investigation. **K.** Data from **I** and **J** presented as a preference for the stressed conspecific as a percentage of total social interaction. Experimental rats in the vehicle condition preferred interaction with stressed juveniles and avoided stressed adults; DCZ eliminated these preferences (ps < 0.006). Atlas images recreated from Paxinos & Watson (1998), use pending permission. Schematic in A created with BioRender.com.

For rats with pIC-PL hM4Di, the same patterns were evident. In SAP tests with juveniles, vehicle treated rats spent more time interacting with the stressed conspecific after vehicle injections, but equal time after DCZ injection (Figure 3I) which was supported by a significant main effect of stress (*F*(1, 8) = 7.417, p = 0.0261, 𝜂^2^ = 0.06), stress by drug interaction (*F*(1, 8) = 7.823, p = 0.0233, 𝜂^2^ = 0.07), and significant post hoc comparison between stress and naive in the vehicle condition (p = 0.0110). In SAP tests with adults, after vehicle treatment rats spent less time interacting with the stressed conspecific and after DCZ treatment time with the conspecific was equal (Figure 3J). This resulted in a significant stress by drug interaction (*F*(1, 10) = 12.06, p = 0.0060, 𝜂^2^ = 0.07) and significant post hoc comparison between naive and stress in the vehicle condition (p = 0.0038). When considering the percent of time spent with the stressed conspecifics, inhibition of pIC input to the PL prevented the preference for stressed juveniles and avoidance of stressed adults (Figure 3K) which resulted in a significant main effect of age (*F*(1, 18) = 8.965, p = 0.003, 𝜂^2^ = 0.14), a significant age by drug interaction (*F*(1, 18) = 23.64, p < 0.001, 𝜂^2^ = 0.32), and significant post hoc comparisons between vehicle and DCZ in juveniles (p = 0.006) and adults (p = 0.005). In summary, inhibition of insular terminals in the PL, regardless of their origin in the pIC or aIC prevented the expression of preference for stressed juveniles and avoidance of stressed adults.

### Prelimbic neurons postsynaptic to insular afferents are necessary for social affective behaviors

Having established that anterior and posterior insula input to the PL is necessary for social affect behavior, we next sought to determine if the PL neurons postsynaptic to that input are also necessary. To inhibit PL_IC_ cells in SAP tests, adult male experimental rats received bilateral injections of an anterograde, transsynaptic viral vector encoding cre-recombinase (AAV1-cre) into the anterior insula or the posterior insula and bilateral injections of a cre-dependent viral vector encoding hM4Di into the PL, allowing for reversible inhibition of the specific population of PL neurons that are postsynaptic to insula. Three weeks later, experimental rats underwent SAP testing with juvenile conspecifics, receiving systemic injections of vehicle (DMSO in saline) or CNO (3 mg/kg, i.p.) 45 minutes prior to testing (Figure 4A). In the rats with AAV1-cre in the aIC, experimental rats preferred interaction with stressed juveniles after vehicle injection but appeared to lose this preference after CNO (Figure 4C), which was supported by a drug by stress interaction (*F*(1,14) = 7.24, p = 0.018, 𝜂^2^ = 0.06), and a significant difference between social investigation of naïve and stressed juveniles in the vehicle condition (p = 0.038) that was not present in the CNO condition (p = 0.461). In SAP tests with adults, experimental rats spent more time investigating naïve adults after vehicle injection but appeared to lose this preference after CNO administration (Figure 4D). We observed a drug by affect interaction (*F*(1,14) = 8.89, p = 0.0099, 𝜂^2^ = 0.07), however, post-hoc comparisons revealed no significant difference in social investigation of naïve and stressed adults in either the vehicle (p = 0.092) or CNO (p = 0.12) condition. When considering the percent of time spent with the stressed conspecifics, inhibition of PL_aIC_ neurons blocked the preference for stressed juveniles and avoidance of stressed adults (Figure 4E). This was supported by a drug by age interaction (*F*(1,14) = 21.8, p = 0.0004, 𝜂^2^ = 0.17) and significant post-hoc comparisons between vehicle and CNO in the juvenile condition (p = 0.015) and adults (p = 0.0073).

**Figure 4.**
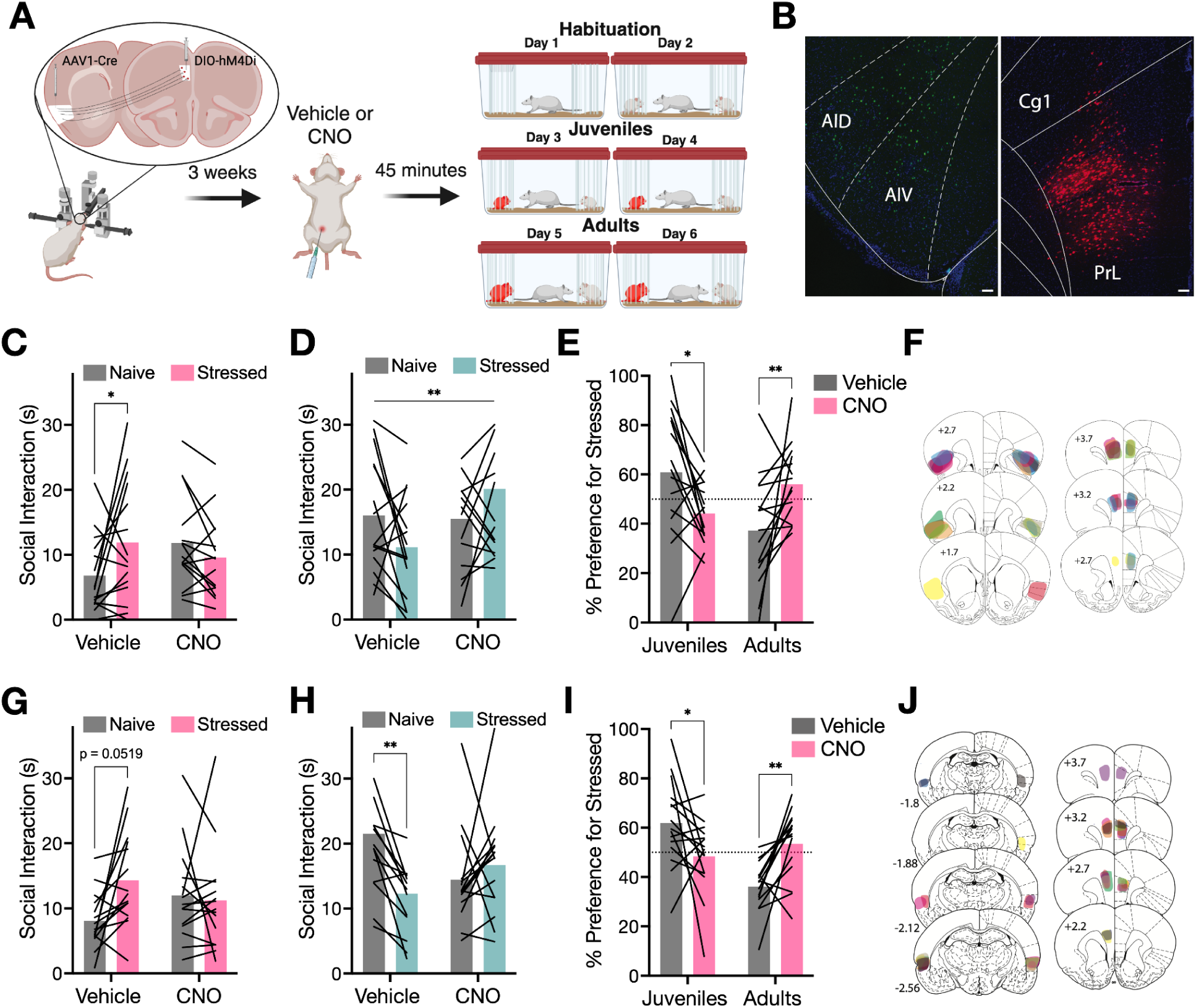
Inhibition of PL_IC_ neurons interferes with social affective behavior. **A.** Schematic diagram of viral injections. Anterograde, transsynaptic AAV1 encoding cre-recombinase was bilaterally injected into the pIC and cre-dependent AAV5 encoding hM4Di was bilaterally injected into the PL. 3 weeks later rats received systemic vehicle (V) or CNO (3 mg/kg) injections 45 min prior to SAP tests with juvenile or adults conspecifics. **B.** (left-to-right): Representative fluorescent images of cre immunoreactivity in the pIC and native DIO-hM4Di-mCherry expression in the PL (green = cre immunoreactivity; red = native mCherry, blue = DAPI; Scale bar = 200 μM). **C.** Mean (with individual replicates) time spent interacting with the naïve and stressed juvenile conspecifics during the 5 min SAP tests after vehicle or CNO injection. In the vehicle condition, rats spent more time investigating stressed juvenile conspecifics than naïve juvenile conspecifics (p = 0.038). In the CNO condition rats explored the stressed and naïve conspecifics equally. **D.** Mean (with individual replicates) time spent interacting with the naïve and stressed adult conspecifics during 5 min SAP tests after vehicle or CNO injection. A drug by social affect interaction effect was observed (p = 0.0099), with CNO injections increasing the time spent investigating the stressed adult (p = 0.118) compared to vehicle injections (p = 0.092). **E.** Data from **C** and **D** expressed as the mean (with individual replicates) preference for the stressed conspecific as a percentage of the total social interaction. CNO injections reduced preference for the stressed juvenile (p = 0.015) and increased preference for the stressed adult (p = 0.007). **F.** Location of aIC cre deposits (left) and PL mCherry expression in the PL (right side, individual rats are represented by different colors) corresponding to data in Panels C-E. **G.** Mean (with individual replicates) time spent interacting with the naïve and stressed juvenile conspecifics during the 5 min SAP tests after vehicle or CNO injection. In the vehicle condition, rats spent more time investigating stressed juvenile conspecifics than naïve juvenile conspecifics (p = 0.052). In the CNO condition rats explored the stressed and naïve conspecifics equally. **H.** Mean (with individual replicates) time spent interacting with the naïve and stressed adult conspecifics during 5 min SAP tests after vehicle or CNO injection. In the vehicle condition, rats spent more time investigating naïve adult conspecifics than stressed adult conspecifics (p = 0.002). In the CNO condition rats explored the stressed and naïve conspecifics equally. **I.** Data from **G** and **H** expressed as the mean (with individual replicates) preference for the stressed conspecific as a percentage of the total social interaction. CNO injections reduced preference for the stressed juvenile (p = 0.016) and naïve adult (p = 0.003). **J.** Location of pIC cre deposits (left) and PL mCherry expression in the PL (right side, individual rats are represented by different colors) corresponding to data in Panels G-I. *p < 0.05, **p < 0.01. Diagram in A was created with BioRender.com. Atlas images recreated from Paxinos & Watson (1998), use pending permission.

In summary, chemogenetic inhibition of PL_aIC_ neurons interfered with social affective preference behaviors toward stressed juveniles and naïve adults.

For inhibition of PL neurons postsynaptic to pIC, experimental rats spent more time interacting with stressed juveniles after vehicle injection but spent equivalent amounts of time investigating both conspecifics after CNO injection (Figure 4G). 2-way ANOVA revealed a main effect of social affect (*F*(1, 14) = 4.298, p = 0.057, 𝜂^2^ =5.75) and a drug by affect interaction effect (*F*(1, 14) = 3.890, p = 0.069, 𝜂^2^ = 6.87) that approached significance. Post-hoc comparison also revealed that the difference in time spent investigating the stressed vs. the naïve juvenile in the vehicle condition approached significance (p = 0.052). In SAP tests with adult conspecifics, experimental rats spent more time investigating naïve adults after vehicle injection but appeared to lose this preference after CNO administration (Figure 4H), which was supported by a drug by stress interaction (*F*(1,14) = 13.1, p = 0.0028, 𝜂^2^ = 0.12). Post-hoc comparison revealed a significant difference between social investigation of naïve and stressed adults in the vehicle condition (p = 0.0021) that was not present in the CNO condition (p = 0.547). Considering the percentage of time spent interacting with the stressed conspecifics, CNO appeared to reverse the patterns that were present after vehicle injections (Figure 4I). We observed a main effect of age (*F*(1,14) = 6.08, p = 0.027, 𝜂^2^ = 0.09) and a drug by age interaction (*F*(1,14) = 24.7, p = 0.0002, 𝜂^2^ = 0.21). Post-hoc comparisons revealed a significant change in preference for stressed juveniles (p = 0.016) and naïve adults (p = 0.0029) when comparing vehicle to CNO. In summary, chemogenetic inhibition of PL_pIC_ neurons interfered with social affective preference behaviors toward stressed juveniles and naïve adults.

### PL_IC_ neurons are primarily glutamatergic regardless of insula origin

To classify PL_IC_ neurons as GABAergic or glutamatergic, tissue from rats with AAV1-cre deposited in either the aIC or pIC were stained for GAD67, a marker for inhibitory neurons, or CaMKII, a marker of excitatory neurons, and imaged under a fluorescence microscope. PL images were taken from both the left and right hemispheres. Cells expressing DAPI, mCherry (PL_IC_ neurons), green fluorescence (GAD67 or CaMKII), and both mCherry and green fluorescence were counted across shallow (Layers II/III) and deep (Layers V/VI) cortical layers (Figure 5A&B). Sections without mCherry expression were excluded. For tissue stained with GAD67, there was a final N of 23 sections from PL_aIC_ tissue and 19 sections from PL_pIC_ tissue that met inclusion criteria for analysis. For tissue stained with CaMKII, 20 sections were obtained from PL_aIC_ tissue and 19 from PL_pIC_ tissue. The percentage of PL_IC_ neurons colabeled with either GAD67 or CaMKII was calculated as a percent of total PL_IC_ neurons. In general, there were more PL neurons that were colabeled with mCherry in rats with AAV1-cre in the aIC, and more found in layers V/VI (Figure 5C). ANOVA revealed main effects of cortical layer (*F*(1,49) = 40.6, p < 0.0001, 𝜂^2^ = 0.09) and insula origin (*F*(1,49) = 46.5, p < 0.0001, 𝜂^2^ = 0.37), and a cortical layer by insula origin interaction effect (*F*(1,49) = 13.3, p = 0.0006, 𝜂^2^ = 0.03). Post-hoc comparisons test revealed significantly more PL_aIC_ neurons in layers V/VI than layer II/III (p < 0.0001) and significantly more PL_aIC_ neurons than PL_pIC_ neurons in layer II/III (p < 0.0001) and layers V/VI (p < 0.0001). Considering cell type, roughly twice as many PL_IC_ neurons colabeled with CaMKII (Figure 5D). A 3-way ANOVA (cell-type x cortical layer x insula origin) revealed a main effect of cell-type (GAD67 vs. CaMKII) (*F*(1,77) = 77.9, p < 0.0001, 𝜂^2^ = 0.39) and both a cell-type by cortical layer interaction (*F*(1,77) = 4.23, p = 0.043, 𝜂^2^ = 0.84) and cell-type by insula origin interaction (*F*(1,77) = 6.65, p = 0.012, 𝜂^2^ = 0.04).

**Figure 5.**
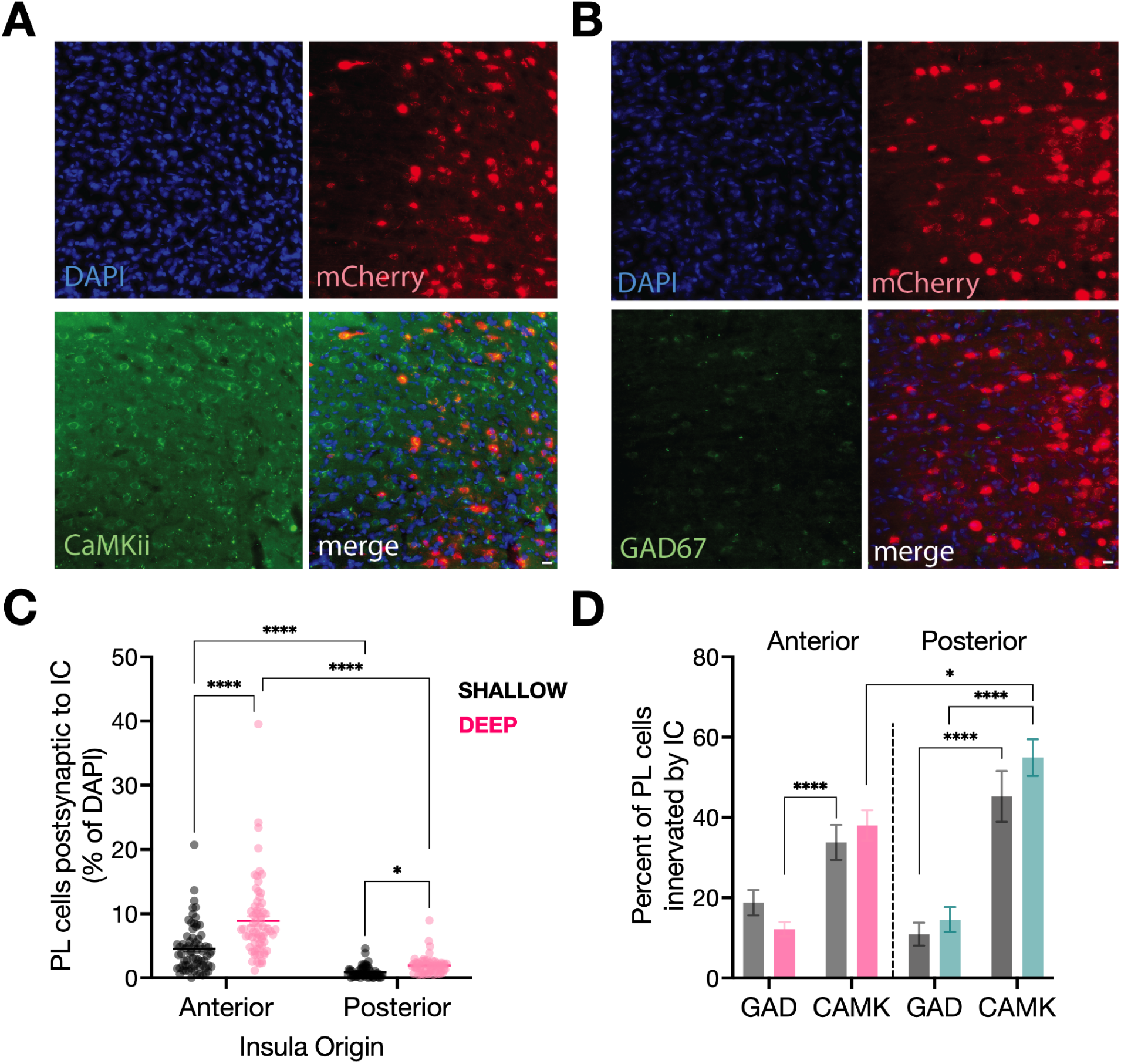
PL_IC_ neurons are primarily glutamatergic. **A.** Representative fluorescent images of nuclei (TOP LEFT, DAPI), PL_IC_ neurons (TOP RIGHT, mCherry), GAD67 neurons (BOTTOM LEFT, GFP), and colabeled PL_IC_ + GAD67 neurons (BOTTOM RIGHT, merge). **B.** Representative fluorescent images of nuclei (TOP LEFT, DAPI), PL_IC_ neurons (TOP RIGHT, mCherry), CaMKii neurons (BOTTOM LEFT, GFP), and colabeled PL_IC_ + CaMKii neurons (BOTTOM RIGHT, merge). Scale bar = 200 μM. **C.** Mean (with individual replicates) percentage of PL that colabeled with mCherry, “PL_IC_ neurons”, within the PL. PL_aIC_ cells represent a larger population of neurons within the PL than PL_pIC_ cells and a larger portion of PL_IC_ cells were found in layers V/VI. **D.** Mean (with SEM) percentage of PL_IC_ cells colabeled with either GAD67 or CaMKii. Significantly more PL_IC_ neurons were colabeled with CaMKii regardless of cortical layer or insula origin. *p < 0.05, ****p < 0.0001.

Post-hoc comparisons test revealed significantly more PL_aIC_ neurons colabeled with CaMII than GAD67 in layers V/VI (p < 0.0001), significantly more PL_pIC_ neurons colabeled with CaMKII than GAD67 in both layers II/III (p < 0.0001) and layers V/VI (p < 0.0001), and significantly more PL_pIC_ neurons than PL_aIC_ neurons colabeled with CaMKii in layers V/VI (p = 0.032). Taken together, PL_IC_ neuronal populations were primarily glutamatergic neurons residing in deep layers, with PL_aIC_ neurons representing a significantly larger population of cells and PL_pIC_ neurons consisting of more glutamatergic neurons relative to PL_aIC_ neuronal populations.

## Discussion

We investigated insula-PL circuit involvement in a social affective preference task in which rats display behaviors that are dependent upon the age and stress experience of the conspecifics. PL inactivation abolished preference for stressed juveniles, indicating a role for this region in social affective decision making. Projections from the insula to the PL arise along the extent of its rostral-caudal axis but are most concentrated in the aIC and pIC. Our prior work established the pIC as critical to approach and avoidance responses to stressed conspecifics but we had not investigated the aIC. Similar to inhibition of the pIC reported previously (Rogers-Carter et al., 2018b, 2019), chemogenetic inhibition of the aIC interfered with social approach to stressed juveniles and avoidance of stressed adults. To test whether the circuit of insula projections and their postsynaptic targets in the PL contributes to social approach and avoidance in the SAP test, we used tract specific chemogenetics to inhibit insula terminals within the PL, and PL neurons that receive monosynaptic input (“PL_IC_ neurons”). Inhibition of afferent axon terminals arising from either aIC or pIC within the PL interfered with both the approach to stressed juveniles and avoidance of stressed adults, suggesting that the insula provides the PL with information necessary for behavior selection. Accordingly, inhibition of PL_IC_ neurons also interfered with social affective preferences. To our knowledge, the current data provide the first mechanistic evidence in rodents implicating the core components of the salience network, insula and prefrontal cortex, to decision-making behaviors. The results enrich our understanding of the neurobiology of social affective decision making by identifying a cortical pathway necessary for approach to or avoidance of stressed others.

The loss-of-function experiments implicate the insula-PL in SAP behaviors, but do not inform us as to what sort of information processing occurs within the circuit.

The SAP test presents a social context where the choice to approach or avoid a conspecific is informed by multiple factors including age, affect, familiarity, and internal stress. Therefore, we explored the question: What aspects of the SAP test recruit the insula-PL circuit? We consider three possibilities. The first is the need for multisensory integration, which we have discussed previously (Rogers-Carter et al., 2018b, 2019; Rogers-Carter and Christianson, 2019). Insula receives input from all sensory modalities including visceral, interoceptive sensation (Allen et al., 1991; Gehrlach et al., 2020) and is capable of multisensory integration as evinced by nonlinear summation of field potentials evoked by multimodal (*e.g.,* somatosensory and auditory) stimulation (Rodgers et al., 2008). Therefore, upon encountering stressed conspecifics, sensory information that provides for interpretation of distress (overt behaviors, vocalizations, chemosignals) and conspecific age (size, chemosignals) may be integrated within the insula and related to the PL. Future investigations using direct recording methods of insula activity will be needed to determine if multisensory integration within the insula during social decision making.

The second possibility concerns the ambiguity inherent in the task. When presented with distressed conspecifics in an otherwise familiar and safe environment, the affective/motivational state of the conspecifics may be ambiguous because the observer is not privy to the cause of the other’s distress. Importantly, there are social scenarios where the motivation for interaction is fundamentally so strong, or innate, that the social behaviors that present may not require the sort of executive functions conventionally associated with the frontal cortex. For example, a rat presented with a predator would behave defensively and a rat presented with an opposite sex conspecific would engage in appetitive, approach behaviors. Although there is ample evidence that insula and prefrontal regions encode features associated reproductive behavior (Kingsbury et al., 2020) and threat/avoidance (Scheggia et al., 2020; Lui et al., 2021), it strikes us as unlikely that primitive escape and approach behaviors to strong social stimuli would require cortical information processing. To test whether the insula-PL circuit is necessary in one such scenario, after the adult and juvenile SAP tests were concluded, we presented rats with two additional tests with a male and female conspecific after vehicle or CNO/DCZ injection. In stark contrast to the findings in social affective discrimination, inhibition of aIC, the insula terminals in the PL, or the PL_IC_ neurons had no effect on preference for female conspecifics (Supplementary Figure S2). Opposite sex preference likely represents one extreme end of a continuum on which social stimuli may be identified as approachable or avoidable. At the extremes, evolutionarily old neural systems generate the reflex-like approach and avoidance behaviors to strong stimuli, while cortical circuits are recruited to select behavior when the social stimuli are more ambiguous, as in the SAP test.

In addition to social and contextual factors, behavior selection is shaped by internal states. For example, in the SAP setting, food deprived rats do not display preferences for stressed juveniles, but those preferences return after a period of feeding (Barretto-de-Souza et al., 2023). Similarly, prior stress treatment of the test rat shifts preferences in adult rats (Toyoshima et al., 2022) and mice (Maltese et al., 2025). It is becoming clear that the insula encodes many aspects of internal states including pain (Jasmin et al., 2004), illness (Koren et al., 2021), hunger, thirst (Livneh et al., 2017), internal temperature (Vestergaard et al., 2023), and fear (Klein et al., 2021) and so it may be the affective component of interaction with stressed conspecifics that recruits the insula-PL circuit. To determine if the insula-PL circuit influences social behavior in the absence of any affective manipulation, we performed a gain-of-function experiment in which rats received an excitatory chemogenetic receptor (hM3Dq) in the aIC or pIC with chemogenetic activation by DCZ in the PL prior to one-on-one social interactions with naive conspecifics (Supplementary Figure S3). Despite this manipulation strongly increasing neural activity in the PL (Supplemental Figure S1), we observed no changes in social interaction. While direct measurement of neural activity in this circuit will be necessary to isolate exactly what features of the internal, social, and contextual stimuli and the ensuing behaviors are encoded in the rat salience network, our additional experiments suggest that ambiguity and interoception are probably reasons this circuit is important to social responses to others in distress.

Our results that the PL is needed for social affective decision making is consistent with its anatomical connections with limbic structures and sensory cortices, and its functional significance to various aspects of social behavior, including social novelty (Zhao et al., 2022), approach (Liang et al., 2018), discrimination of affective states in rodents (Scheggia et al., 2020), and short-term recognition memory (Yashima et al., 2023). Numerous *in vivo* electrophysiology and calcium imaging studies reveal that the mPFC dynamically tracks changes in social environments (Zhao et al., 2022), encodes various social sensory cues (Levy et al., 2019), and modulates activity in anticipation of potential threats (Kim et al., 2018), indicating that this region is responsible for updating representations of social contexts. The range of social factors that appear to engage the mPFC might be due to the intrinsic mixed selectivity that allows the mPFC activity to be dynamically reorganized as contexts, motivations, or behavioral options change (Rigotti et al., 2013). Thus, all sorts of social interactions should have prefrontal neural correlates, regardless of whether these correlates are necessary for behavior.

Here, PL_IC_ neurons were primarily glutamatergic, regardless of insula origin or cortical layer. These neurons include a mix of intratelencephalic cells projecting to cortical, striatal, claustral, and amygdalar regions, pyramidal tract cells that target subcortical regions, and corticothalamic cells which project to thalamic relay nuclei and the thalamic reticular nucleus (Anastasiades and Carter, 2021), suggesting that insula input to the PL could influence sensory, motivational and motor processes via afferents to the BLA, dorsal raphe nucleus, and periaqueductal gray which bidirectionally modulate social preference, induce active escape-like behavior, and promote social interaction, respectively (Warden et al., 2012; Franklin et al., 2017; Huang et al., 2020). Future studies will focus on PL projections to the nucleus accumbens and amygdala, which shape approach or avoidance, respectively (Murugan et al., 2017; Diehl et al., 2020).

To our knowledge, this is the first causal demonstration the two key nodes of the salience network, insula and PL, are required for affective behavioral decision making which reinforces numerous functional neuroimaging experiments that correlate social dysfunction to the salience network (Nagai et al., 2007; Uddin et al., 2013; Goodkind et al., 2015; Uddin, 2015; Namkung et al., 2017). However, we examined the insula-PL circuit from only one direction. Future studies must investigate how the PL may shape insula encoding through its reciprocal connections (Shi and Cassell, 1998). Although we examined how the salience network is engaged in the context of social dilemmas, we argue here that it is some aspect of the multisensory integration and ability to organize situationally-relevant factors like ambiguity and internal states that necessitates cortical involvement. We expect that our findings will generalize to non-social domains and inform a theory of insula-prefrontal function that accounts for the growing array of behaviors and cognitive processes associated with these regions.

## Acknowledgements

The authors wish to thank Dr. Bret Judson, director of the Boston College Core Imaging Facility; Nancy McGilloway and Todd Gaines, administrators of the Boston College Animal Care Facility; and Zenia Bathena, Elizabeth Schwartz, Christopher Catalano, Ricardo Mora, Natalie Cortopasi and Joshua Elbaz for their contributions to data collection. This project was funded by a grant from the National Institutes of Health (MH119422) and Undergraduate Research Fellowships from Boston College.

## Author Contributions

Conception: EC, AD, MRM, JPC; Data Collection: EC, AD, TM, ZB, AN; Data Analysis: EC, AD, TM, JPC; Manuscript Preparation: EC, AD, JPC; Final Editing: EC, JPC; Funding Acquisition: JPC.

## Supplemental Materials

### Verification of chemogenetic function with Fos analysis

4 days after SAP testing, rats from the chemogenetic manipulation studies underwent one-on-one social interaction tests to elicit c-Fos activation in the regions of interest. Here we sought to confirm that DCZ inhibited aIC or PL activity as intended by comparing c-Fos activation between experimental rats that received vehicle vs. DCZ injections. Importantly, we previously provided the same sort of verification for CNO using the same viral strategy employed here (Rogers-Carter et al., 2018a; Djerdjaj et al., 2022). Each experimental rat was placed into a standard plastic tub cage with beta chip bedding and a wire lid 1 h prior to testing. Experimental rats then received vehicle or drug injections 45 minutes prior to testing. Testing consisted of a stressed juvenile being introduced into the experimental rat’s cage for 5 minutes and returned to the home cage. 90 minutes after testing, rats were perfused and brains were collected and sectioned as described above. To verify inhibition of the PL or aIC via DCZ administration, tissue sections were washed in PBS-T (0.01% Triton-X 100), blocked in 5% normal donkey serum in PBS-T, and then incubated overnight in rabbit polyclonal antibody to c-fos (1:5,000; EnCor RPCA-c-Fos-AP, Lot #:241-102320). Sections were then washed in PBS-T and incubated in AlexaFluor 488 AffiniPure donkey anti-mouse fluorescent secondary antibody (1:500; Jackson Immunoresearch, Cat #715-545-150). Sections were then floated onto glass slides and coverslipped with Vectashield containing DAPI. The PL was imaged as described above and for each image the total number of PL_aIC_, c-Fos, and PL_aIC_ cells colocalized with c-Fos were counted.

**Supplemental Figure S1.**
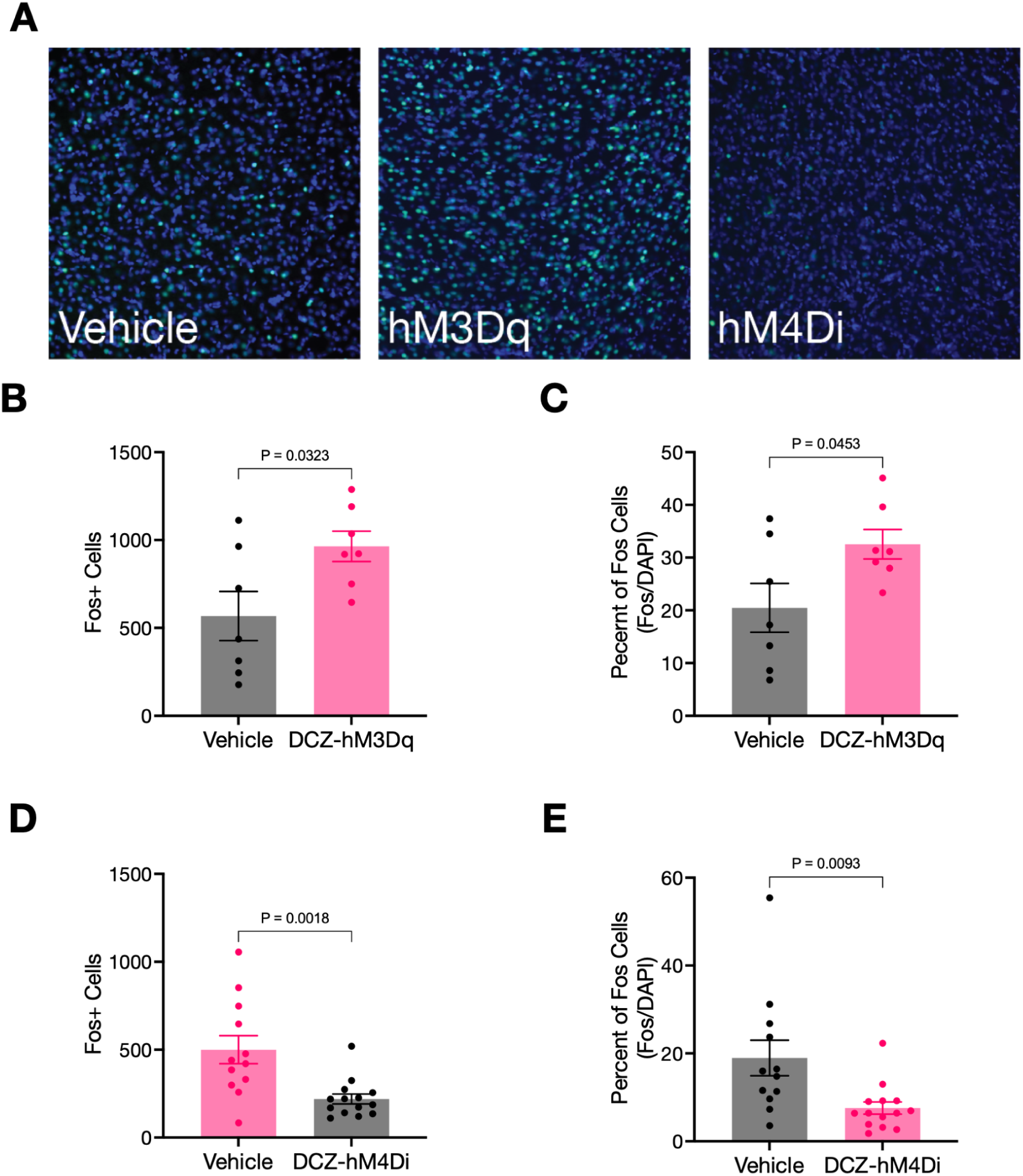
Verification of DCZ in the PL. **A**. Representative Fos (green) and DAPI (blue) immunofluorescence from PL of rats treated with vehicle or DCZ with either hM3Dq or hM4Di. **B.** Mean (+/- SEM) number of Fos cells in rats with hM3Dq in the insula and DCZ in the PL. **C.** Data in B expressed as a percentage of total cells (DAPI). **D**. Mean (+/- SEM) number of Fos cells in rats with hM4Di in the insula and DCZ in the PL. **C**. Data in D expressed as a percentage of total cells (DAPI). P-values denote the significance of independent samples t-tests.

### Effect of insula-prefrontal inhibition on opposite sex conspecific preference

To determine if the insula-prefrontal circuits under study contribute specifically to social affective behaviors, or if they are involved in sociability/preference more generally, after the juvenile and adult versions of the SAP tests were complete, rats were given 2 additional tests which were identical in every way to the SAP tests with stressed conspecifics, except in this case the conspecifics were an unfamiliar, naive to stress male and an unfamiliar, naive to stress female (Supplemental Figure S2A). Adult male test rats exhibit a robust preference for interaction with adult females (Supplemental Figure S2B-E). As with the SAP tests, all rats were tested after vehicle injection on one day, and after CNO or DCZ on the other day with test order counter balanced.

**Supplemental Figure S2.**
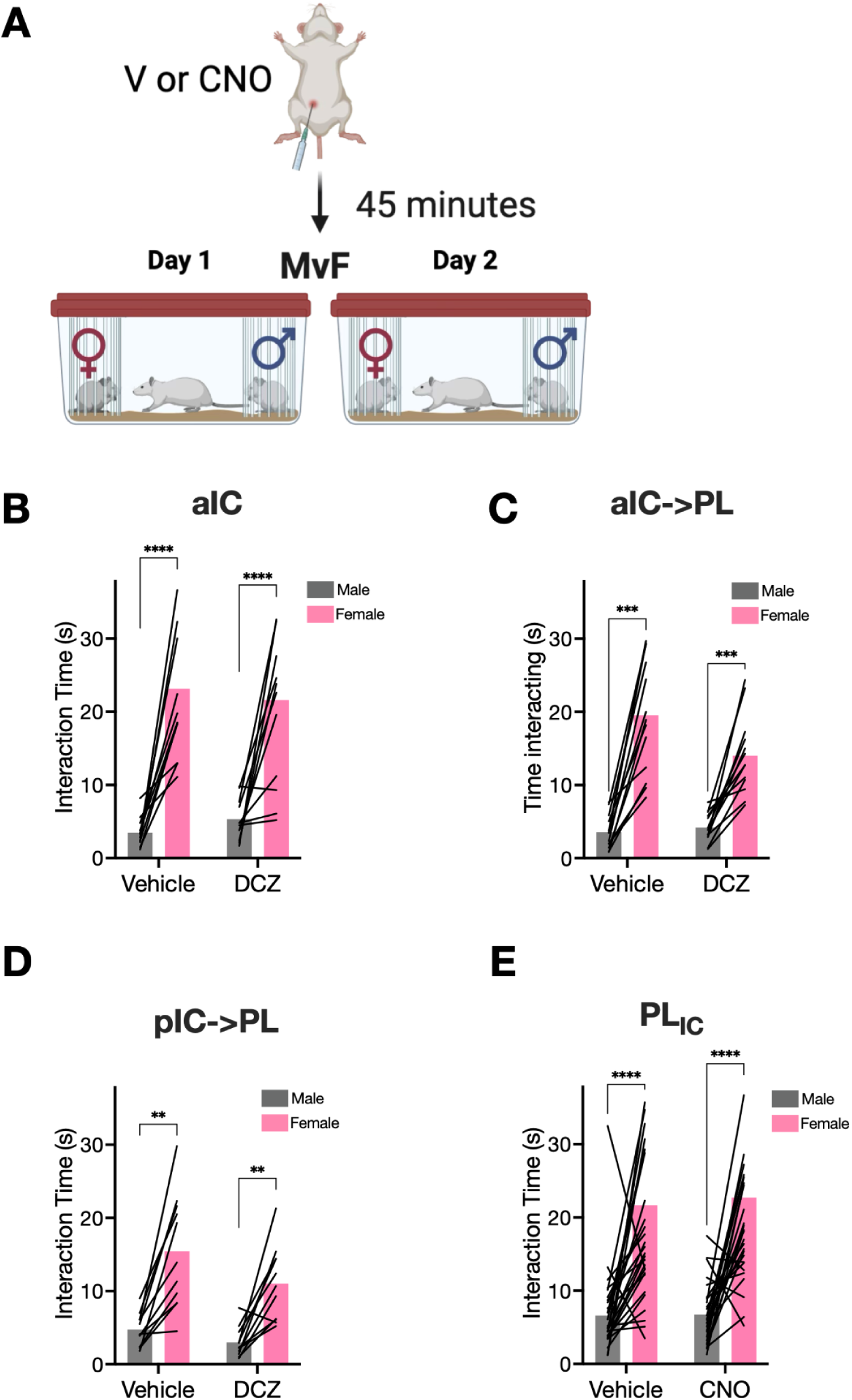
Inhibition of insula input to the PL does not alter opposite sex preference. **A.** Schematic of opposite sex preference tests. **B-E.** Mean with individual replicates, time spent interacting with male or female conspecifics after DCZ or CNO treatment in rats with hM4Di in the aIC (**B**) from experiments described in Figure 2, at the aIC-PL terminals (**C**), at the pIC-PL terminals (**D**) from experiments described in Figure 3, or in PL neurons postsynaptic to insula input (**E**). Data in E include both pIC and aIC versions of the experiment described in Figure 4 of the main text. In all cases there were significant main effects of sex (ps< 0.001), but no interactions or main effects of drug. Asterisks indicate significance of post hoc Sidak comparisons (*p < 0.05, **p<0.01, ***, p<0.001, ****p< 0.0001).

### Chemogenetic stimulation of insula terminals in the PL

Because the results of the inhibitory chemogenetic experiments present a compelling case that the pIC and aIC input to the PL are necessary for social affective decision making here we sought to establish whether stimulation of these tracts were sufficient to alter social investigation behavior in the absence of any affective manipulation. We previously demonstrated that modulation of the insula by oxytocin (Rogers-Carter et al., 2018b) or by corticotrophin releasing factor (Rieger et al., 2022a) could induce social approach to naive juveniles and reduce social interaction with naive adults. Here we made deposits of the Gq coupled chemogenetic receptor hM3Dq, (AAV5-hSyn-hM3Dq-mCherry; Addgene Cat. No. 50474-AAV5), in either aIC or pIC with cannulas placed in the PL exactly as described for the terminal inhibition experiments described in Figure 3. Rats were given 5 minute social interaction tests with an unfamiliar juvenile conspecific in a standard tub cage as the test arena. DCZ or vehicle was microinfused to the PL 45 min prior to test; drug order was counterbalanced and DCZ efficacy was established with Fos (Supplemental Figure S2). If stimulating the insula input to the PL is sufficient to alter social behavior itself, then DCZ would alter social interactions regardless of stress. However, we found that DCZ had no effect on social interaction in either aIC or pIC variations of this experiment (Supplemental Figure S3).

**Supplemental Figure S3.**
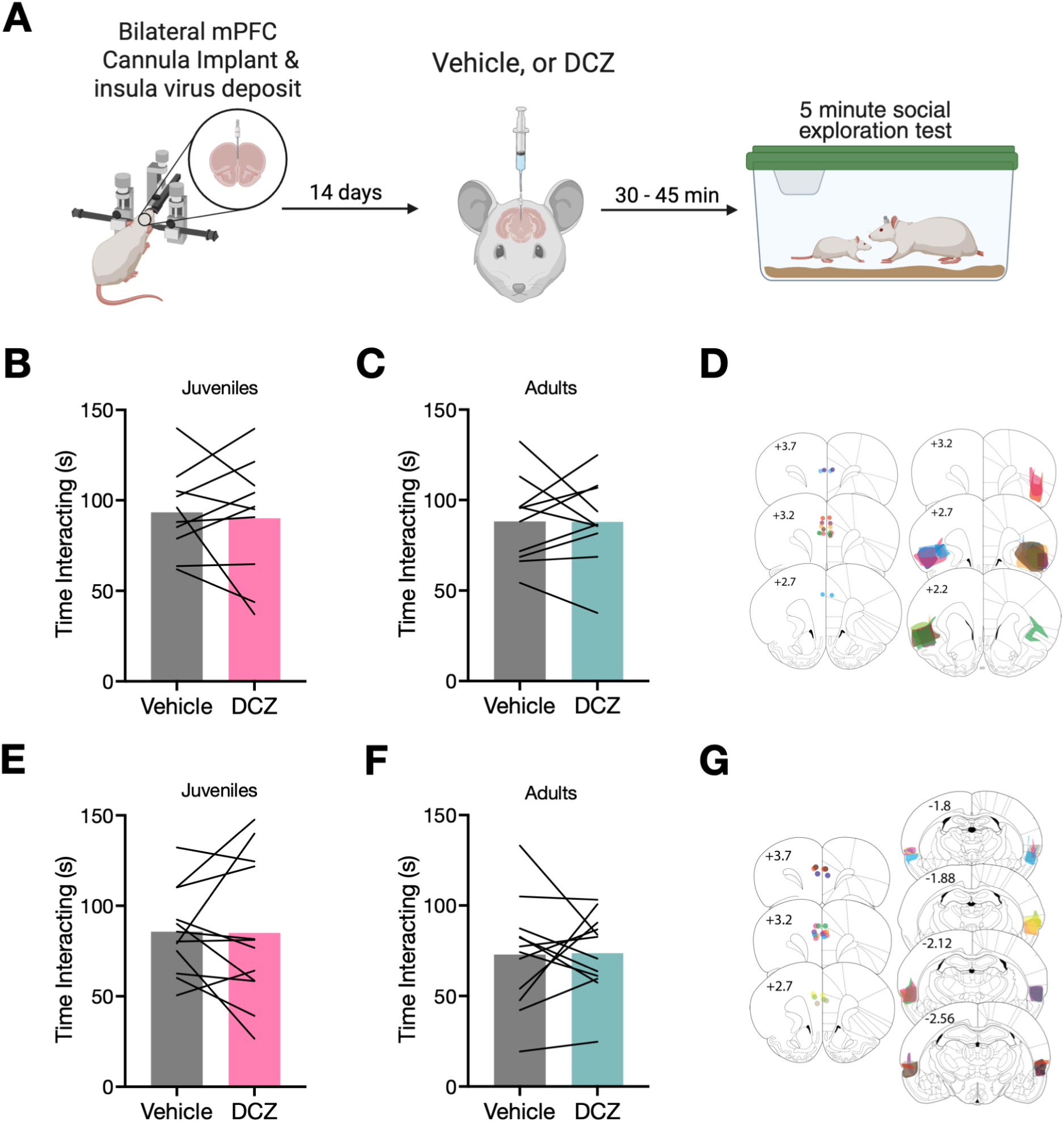
Chemogenetic stimulation of insula terminals in the PL. **A.** Schematic of approach. **B-C.** Mean (with individual replicates) time spent interacting with naive juvenile (B) or adult (C) conspecifics in rats with hM3Dq in the aIC and cannula in the PL. **D.** Representative location of PL cannula and aIC virus positions. **E-F.** Mean (with individual replicates) time spent interacting with naive juvenile (E) or adult (F) conspecifics in rats with hM3Dq in the pIC and cannula in the PL. **D.** Representative location of PL cannula and pIC virus positions. No significant effects were observed in any experiment.

## Notes

### Competing Interest Statement

The authors have declared no competing interest.

